# Brief sensory deprivation triggers plasticity of dopamine-synthesising enzyme expression in genetically labelled olfactory bulb dopaminergic neurons

**DOI:** 10.1101/2020.06.03.132555

**Authors:** Darren J Byrne, Marcela Lipovsek, Andres Crespo, Matthew S Grubb

**Affiliations:** Centre for Developmental Neurobiology, Institute of Psychiatry, Psychology and Neuroscience (IoPPN), King’s College London, London, SE1 1UL, UK; Ear Institute, University College London, 332 Gray’s Inn Road, London, WC1X 8EE, UK

## Abstract

In the glomerular layer of the olfactory bulb, local dopaminergic interneurons play a key role in regulating the flow of sensory information from nose to cortex. These dual dopamine- and GABA-releasing cells are capable of marked experience-dependent changes in the expression of neurotransmitter-synthesising enzymes, including tyrosine hydroxylase (TH). However, such plasticity has most commonly been studied in cell populations identified by their expression of the enzyme being studied, and after long periods of sensory deprivation. Here, instead, we used brief 1- or 3-day manipulations of olfactory experience in juvenile mice, coupled with a conditional genetic approach that labelled neurons contingent upon their expression of the dopamine transporter (DAT-tdTomato). This enabled us to evaluate the potential for rapid changes in neurotransmitter-synthesising enzyme expression in an independently identified neuronal population. Our labelling strategy showed good specificity for olfactory bulb dopaminergic neurons, whilst revealing a minority sub-population of non-dopaminergic DAT-tdTomato cells that expressed the calcium-binding protein calretinin. Crucially, the proportions of these neuronal subtypes were not affected by brief alterations in sensory experience. Short-term olfactory manipulations also produced no significant changes in immunofluorescence or whole-bulb mRNA for the GABA-synthesising enzyme GAD67/*Gad1*. However, in bulbar DAT-tdTomato neurons brief sensory deprivation was accompanied by a transient, small drop in immunofluorescence for the dopamine-synthesising enzyme dopa decarboxylase (DDC), and a sustained decrease for TH. Deprivation also produced a sustained decrease in whole-bulb *Th* mRNA. Careful characterisation of an independently identified, genetically labelled neuronal population therefore enabled us to uncover rapid experience-dependent changes in dopamine-synthesising enzyme expression.

## Introduction

Dopaminergic neurons in the mammalian olfactory bulb form a neurochemically-distinct population of glomerular layer interneurons (Hökfelt *et al*., 1975; Kosaka & Kosaka, 2007, 2011). They are dual-transmitter cells, releasing both dopamine and GABA to locally modulate information being sent from nose to cortex (Borisovska *et al*., 2013; Liu *et al*., 2013, 2016; Banerjee *et al*., 2015; Pignatelli & Belluzzi, 2017; Vaaga *et al*., 2017). Production of these transmitters is undertaken by specialised enzymatic pathways, which include tyrosine hydroxylase (TH) as the rate-limiting step in dopamine synthesis, and DOPA decarboxylase (DDC) as its immediate downstream partner (Halász *et al*., 1977). Glutamate decarboxylase (GAD)-67, on the other hand, is the GABA-synthesising enzyme that is ubiquitously expressed in this neuronal class (Kiyokage *et al*., 2010; Kosaka *et al*., 2019).

Expression levels of these neurotransmitter-synthesising enzymes are a well characterised target of activity-dependent plasticity in olfactory bulb dopaminergic neurons. TH, especially, is exquisitely sensitive to changes in sensory input. Since the landmark descriptions of decreased TH enzymatic activity and immunofluorescence in the bulb following damage to the peripheral olfactory system (Nadi *et al*., 1981; Baker *et al*., 1983), a wealth of evidence has shown that both *Th* mRNA and TH protein levels in bulbar cells track alterations in olfactory sensory neuron activity (Baker *et al*., 1984, 1993, 1999; Kosaka *et al*., 1987; Baker, 1990; Stone *et al*., 1990; Cho *et al*., 1996; Philpot *et al*., 1998; Saino-Saito *et al*., 2004; Bastien-Dionne *et al*., 2010; Cave *et al*., 2010; Weiss *et al*., 2011; Grier *et al*., 2016). Indeed, these TH changes are so reliable and sensitive that they are often used as confirmation that particular olfactory system manipulations have been effective (Wilson & Sullivan, 1995; Mandairon *et al*., 2006; Kass *et al*., 2013). Olfactory bulb GAD67 has received rather less attention, but nevertheless has also been shown to undergo experience-dependent changes in both mRNA (*Gad1*) and protein levels (Parrish-Aungst *et al*., 2011; Lau & Murthy, 2012; Banerjee *et al*., 2013; Wang *et al*., 2017).

The vast majority of these activity-dependent enzyme expression changes have been detected following relatively prolonged manipulations of sensory input, usually on the order of weeks. However, there are indications that transmitter-synthesising enzyme expression can change more rapidly. A significant decrease in whole-bulb *Th* mRNA has been observed after only 2 d of unilateral naris occlusion (Cho *et al*., 1996). *In vitro*, increases in TH immunofluorescence or TH-GFP transgene levels have been observed after just 24 h of sustained depolarisation in dissociated and slice cultures, respectively (Akiba *et al*., 2007; Chand *et al*., 2015). We also recently found that a 24 h period of *in vivo* sensory deprivation was associated with a decrease in relative immunofluorescence in olfactory bulb TH-positive neurons, without any accompanying change in their density within the glomerular layer (Galliano et al., 2021).

A significant caveat is that, to date, these changes in neurotransmitter-synthesising enzyme expression have all been detected either by employing tissue-level analyses that encompass the full range of bulbar cell types, or by using the measured variable itself (e.g. TH immunofluorescence) to identify individual cells to be studied. Ideally, gaining cell-type-specific information regarding plasticity in enzyme expression would instead use an independent marker of cell identity. This would enable experience-dependent changes to be identified amongst a consistently identified group of neurons, without the potential distortion that can result when a plastic variable determines which cells to measure. Here, we therefore took advantage of a conditional mouse transgenic line, DAT^IRES*cre*^ (Bäckman *et al*., 2006), which when crossed with an appropriate reporter line generates selective fluorescent label in cells that express the dopamine transporter (DAT-tdTomato). We quantitatively characterised the selectivity of this line and found that it labels a small subset of calretinin-expressing cells in addition to a majority of olfactory bulb dopaminergic neurons. Crucially, this subtype specificity was unaffected by manipulations used to induce brief olfactory deprivation. We found that such short-term, naturally relevant sensory manipulations do not affect relative levels of GAD67 immunofluorescence in olfactory bulb DAT-tdTomato cells, nor whole-bulb levels of *Gad1* mRNA. In contrast, deprivation is associated with decreased whole-bulb *Th* mRNA levels. It also leads to small transient and larger sustained decreases in relative immunofluorescence for DDC and TH, respectively, within an independently identified population of DAT-tdTomato neurons.

## Methods

### Animals

A total of 69 mice of either sex were used. Mice were housed under a 12 h light:dark cycle in an environmentally controlled room with access to water and food ad libitum. Day of birth was designated as P0. DAT^IRES*cre*^ mice (B6.SJL-*Slc6a3*^*tm1*.*1(cre)Bkmn/J*^, Jax stock 006660; Bäckman *et al*., 2006) were crossed with Rosa26-floxed stop tdTomato reporter mice (Gt(ROSA)26Sor^tm9(CAG-tdTomato)Hze^, Jax stock 007909) to generate DAT-tdTomato mice. Wild type C57/Bl6 mice (Charles River, Harlow, UK) were used to back cross each generation of transgenic mice. All experiments were performed at King’s College London under the auspices of UK Home Office personal and project licences held by the authors.

### Unilateral naris occlusion

Olfactory sensory deprivation was induced by unilateral naris occlusion, using a custom made plug (Cummings *et al*., 1997). Each plug was constructed by knotting suture (Ethilon Monofilament Polyamide 6; Ethicon, Cincinnati, USA) around a piece of unscented dental floss pulled through the lumen of PTFE tubing with an outer diameter of 0.6 mm and inner diameter of 0.3 mm (VWR International, cat#: S1810 04). Mice were briefly anaesthetised with isoflurane and the plug trimmed, coated in petroleum jelly and inserted into the right naris. Mice were returned to their home cage and left for 1 or 3 days before tissue collection. Presence of the nose plug was confirmed by examining the nasal cavity after olfactory bulb dissection *ex vivo*. Sham controls, used for qPCR experiments only, had a nose plug inserted into the right naris and immediately removed. Anaesthetised controls, used for all other experiments, were handled and anaesthetised briefly with isoflurane but had no plug inserted into either naris.

### Transcardial perfusion and histology

P28 and P30 DAT-tdTomato mice were anaesthetised with an intraperitoneal overdose of pentobarbital, then transcardially perfused with 20 mL phosphate buffered saline (PBS) with 20 U/mL heparin, followed by 20 mL of 4 % paraformaldehyde (PFA; TAAB Laboratories, Berkshire, UK) in PIPES buffer (3 % sucrose, 60 mM PIPES, 25 mM HEPES, 5 mM EGTA, and 1 mM MgCl_2_; unless otherwise stated, all reagents were from Sigma). PFA was prepared in batches, frozen at -20 °C and defrosted shortly before use. On completion of perfusion, the mouse was decapitated, and the brain removed from the skull and post-fixed in 4% PFA overnight. The brain was then washed with PBS (3x5 minutes) and stored in PBS with 0.05% sodium azide for up to 6 months. For sectioning, brains were embedded in 6% agarose and sliced coronally at 50 μm using a vibratome (Leica VT1000s; Wetzlar, Germany). Slices were stored in PBS and 0.05% sodium azide at 4 °C before staining.

### Immunohistochemistry

Immunohistochemical labelling was always performed on 3 slices from each mouse, at approximately 25%, 50% and 75% of the distance along the anteroposterior axis of the olfactory bulb. Free-floating slices were incubated at room temperature (RT; 18-24 °C) for 2 hours in a blocking and permeabilising solution (10% normal goat serum in PBS containing 0.25% Triton-X and 0.02% sodium azide). They were then incubated in primary antibody diluted in the blocking solution at 4 °C for between 18-24 hours (see Table 1 for antibodies used and their working dilution; endogenous tdTomato fluorescence was always enhanced with immunolabel for red fluorescent protein; mouse anti-TH antibody was used for all staining except when TH was co-labelled with DDC in Fig.1B, in which case rabbit anti-TH was used to avoid host species cross reactivity). For immunofluorescence intensity analysis, slices were incubated for approximately 72 hours in primary antibody solution. Slices were then washed with PBS (3x5 minutes) and incubated in the species-appropriate secondary antibody (Alexa Fluor® Life Technologies) diluted 1: 1,000 in the blocking solution at RT for 2 hours. After 3x5 minutes washes in PBS, slices were mounted on glass slides with MOWIOL® (Millipore).

**Table 1:** Details of primary antibodies

| Antigen | Host Species | Working Dilution | Supplier (product code) |
| --- | --- | --- | --- |
| Calbindin | Rabbit | 1: 5,000 | Swant (CB38a) |
| Calretinin | Mouse | 1: 5,000 | Millipore (MAB1568) |
|  | Rabbit | 1: 5,000 | Swant (7699/3H) |
| cFos | Mouse | 1: 1,000 | Santa Cruz (sc166940) |
| Dopa decarboxylase (DDC) | Mouse | 1: 1,000 | Santa Cruz (sc-293287) |
| Glutamate decarboxylase 67 (Gad67) | Mouse | 1: 500 | Millipore (MAB351R) |
| Red fluorescent protein (tdTomato) | Rabbit | 1: 1,000 | Clontech (632496) |
|  | Rat | 1: 1,000 | Chromotek (5F8) |
| Tyrosine hydroxylase (TH) | Mouse | 1: 1,000 | Millipore (MAB318) |
|  | Rabbit | 1: 1,000 | Millipore (AB152) |

**Figure 1.**
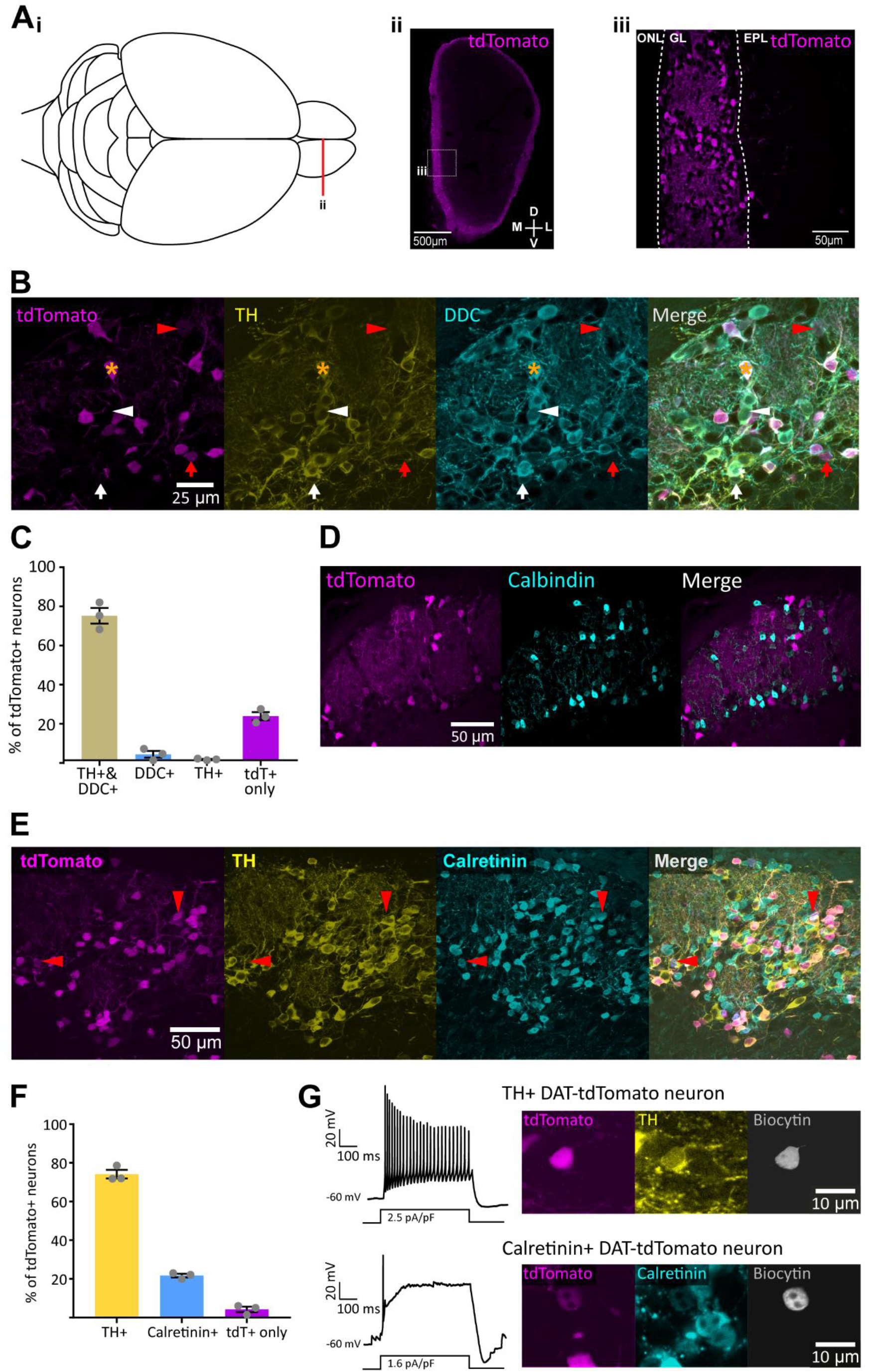
Cell-type identity of olfactory bulb DAT-tdTomato neurons. **A)** tdTomato label in DAT-tdTomato olfactory bulb; **i)** aerial view diagram of mouse brain, showing location of olfactory bulb coronal section in **ii**; **ii)** coronal section of olfactory bulb from a DAT-tdTomato mouse; box shows location of detail in **iii**; **iii)** tdTomato-positive neurons are located predominantly in the olfactory bulb’s glomerular layer (GL); ONL, olfactory nerve layer; EPL, external plexiform layer. **B)** Example maximum intensity projection images from the glomerular layer in the olfactory bulb of a control DAT-tdTomato mouse, showing co-label for TH and DDC. Star indicates location of a neuron showing expression of all three markers. Arrowheads indicate location of neurons expressing two markers: red = DAT-tdTomato and DDC; white = TH and DDC. Arrows indicate location of neurons expressing a single marker: red = DAT-tdTomato; white = DDC. **C)** Mean ± SEM percentage of tdTomato+ neurons positive for TH and/or DDC. Grey circles indicate percentages from individual mice. **D)** Example maximum intensity projection images showing immunofluorescence for calbindin in the olfactory bulb glomerular layer of a control DAT-tdTomato mouse. **E)** Example maximum intensity projection images showing immunofluorescence for TH and calretinin in the glomerular layer of the olfactory bulb in a control DAT-tdTomato mouse. Red arrowheads indicate example neurons expressing DAT-tdTomato and calretinin, but not TH. **F)** Mean ± SEM percentage of tdTomato+ neurons expressing TH and calretinin. Grey circles indicate percentages from individual mice. **G)** Action potential firing patterns in response to 500 ms somatic current injection (left) and immunohistochemical characterisation (right) in example biocytin-filled DAT-tdTomato neurons that were co-labelled for either TH (top) or calretinin (bottom).

### Image acquisition and experimental design

All images were acquired with a laser scanning confocal microscope (Zeiss LSM 710 Axio Examiner Z1) using appropriate excitation and emission filters, a pinhole of 1 AU and a 40x oil immersion objective. Laser power, gain and offset were manually set to prevent fluorescence saturation and ensure optimum signal-to-noise.

For analyses of relative immunofluorescence intensity, we aimed to minimise and control for between-section experimental variability. To reduce the effect of the homecage environment, each experimental ‘set’ of four mice belonged to the same litter and was exposed to the same external odour environment until unilateral naris occlusion or control procedure at P27. All mice were returned to the homecage and left for 1 or 3 days. Perfusions and post-fixing were performed using the same batch of PFA for each set, and mice within each set were perfused within 30 minutes of each other. Brains from the same set were trimmed to the same size and co-embedded in 6% agarose to ensure all brains would be sliced together. The agarose block was cut with landmarks to ensure each mouse could be reliably identified. Co-embedded slices were processed for immunohistochemistry as described above, with primary antibodies applied for 72 hours. Therefore, all control and occluded sections on each slice were exposed to identical reagents at identical concentrations for identical periods of time. The olfactory bulb sections in a co-embedded slice were then imaged on the same day using identical laser and detection settings that were optimised for that co-embedded slice. Z-stacks were taken from different parts of an olfactory bulb slice, setting the start and end Z positions using the DAT-tdTomato fluorescence. Images were acquired at a resolution of 512 × 512 pixels (0.277 μm /pixel) and a Z step of 1 μm.

Images for analysing colocalisation of DAT-tdTomato with other markers were acquired at a resolution of 512 × 512 pixels (0.277 μm /pixel) and a Z step of 1 μm, with multiple Z stacks taken around the entire olfactory bulb section.

All immunohistochemistry, imaging and image analysis of tissue from mice that had been manipulated was performed blind to experimental group.

### Image analysis

All image analysis was performed using ImageJ software (Fiji). Cells were counted manually using the cell counter plugin and only if their soma was entirely inside the image area. Soma size and fluorescence intensity were measured at the neuron’s widest area in the Z-stack, identified when the soma was larger than in the slice above and below. For measuring immunofluorescence intensity, neurons were manually identified for analysis solely by the expression of DAT-tdTomato. TH and GAD67 immunolabel were located primarily outside the nucleus, therefore, average immunofluorescence intensity (mean grey value; MGV) was measured in the somatic cytoplasm only, using a hand-drawn region of interest (ROI). For DDC and cFos, MGV was measured across the entire soma, again using a hand-drawn ROI. All tdTomato cells that had the entire cell body within the image area and for which the widest soma area could be confirmed were measured in each stack.

For immunofluorescence intensity analysis, background MGV values were taken from unlabelled bulbar regions of each Z-stack and were subtracted from the MGV of each cell in that Z-stack. Any resulting negative values (i.e. where background > cell) were set to zero. Normalised percentage immunofluorescence intensity values were then obtained for each cell, by dividing that cell’s background-subtracted MGV value by the combined control average in each slice (over both 1 day and 3 day sections), multiplied by 100.

### Electrophysiology

For acute slice preparation, mice were deeply anaesthetised with isoflurane and decapitated. The olfactory bulb and frontal cortices were rapidly removed and transferred into ice-cold sucrose slicing medium (containing in mM: 240 sucrose, 5 KCl, 1.25 Na_2_HPO_4_, 2 MgSO_4_, 1 CaCl_2_, 26 NaHCO_3_ and 10 D-glucose, bubbled with 95% O_2_ °and 5% CO_2_). 300 μm horizontal sections of the olfactory bulb were cut using a Leica vibratome (VT1000S; Wetzlar, Germany) and were transferred to standard artificial CSF (ACSF; containing in mM: 124 NaCl, 5 KCl, 1.25 Na_2_HPO_4_, 2 MgSO_4_, 2 CaCl_2_, 26 NaHCO_3_ and 20 D-glucose; with 10 μM NBQX, 50 μM APV, 10 μM SR-95531 to block major ionotropic synaptic receptors, continuously bubbled with 95% O_2_ and 5% CO_2_, pH 7.4, ∼310 mOsm). Slices were left to recover at RT for at least 45 minutes before experiments began.

For whole-cell recordings, individual slices were secured in a chamber mounted on a Nikon FN1 microscope. DAT-tdTomato neurons were visualised with a 40x water immersion objective and tdTomato fluorescence was revealed by under LED illumination (pE-100; CoolLED, Andover, UK) with appropriate excitation and emission filters (ET575/50m, CAIRN Research, Kent, UK). Slices were continuously superfused with ACSF heated to physiologically relevant temperature (34 ± 1 C) using an in-line solution heater (TC-344B; Warner Instruments, Kent, UK). Recordings were carried out using a EPC10/2 amplifier coupled to PatchMaster acquisition software (Heka Elektronik, Reutlingen, Germany). Signals were digitised and sampled at 20 kHz. Recording electrodes were pulled from borosilicate glass [outer diameter (OD), 1.5 mm; inner diameter (ID), 1.17 mm] (GT100T-10; Harvard Apparatus, Cambridge, UK) with a vertical puller (PC-10; Narishige, London, UK) and the tip fire-polished with a microforge (MF-830; Narishige, London, UK). Pipettes with 3-5 MΩ resistance were filled with K-gluconate-based intracellular solution, containing, in mM: 124 K-gluconate, 9 KCl, 10 KOH, 4 NaCl, 10 HEPES, 28.5 Sucrose, 4 Na_2_ATP, 0.4 Na_3_GTP, plus 0.15 % biocytin; pH ∼7.3, 280-290 mOsm. To study multiple action potential firing, 500 ms somatic current injections of Δ+5 pA were applied from a holding potential of -60 mV in current-clamp mode with 100 % bridge balance, until the point where the neuron entered depolarisation block.

After completion of electrophysiology experiments, the pipette was gradually removed with a small amount of positive pressure applied to try and dislodge the patched neuron from the pipette. Once the pipette was separated from the cell, the acute slice was transferred to 4% PFA. The slice was fixed overnight at 4° C, washed with PBS (3x5 minutes) and stored in PBS and 0.05 % sodium azide at 4° C, for up to 3 months. Standard immunohistochemistry protocols were used (see above). Streptavidin 488 (Alexa Fluor® Life Technologies) diluted 1: 1,000 was included with the secondary antibodies to label biocytin in the patched neuron.

### Quantitative PCR

All experimental steps for qPCR were performed in compliance with the MIQE guidelines (Bustin *et al*., 2009). One or 3 days after naris occlusion or control sham surgery, wild-type mice were sacrificed by cervical dislocation followed by decapitation. The brain was quickly removed and placed in cold PBS. The right olfactory bulb was dissected and placed in RNAlater (Thermo Fisher Scientific) overnight at 4° C and stored at -20° C until further processing. Total RNA extraction was performed using the RNAqueous® Micro Kit (Life Technologies). RNA was eluted in 20 µl of elusion buffer and quantified using Nanodrop (Thermo Fisher Scientific) and Qubit (RNA HS Assay Kit, Life Technologies). 300-500 ng of total RNA were used for cDNA synthesis with SuperScript™ II Reverse Transcriptase (Life Technologies), 0.5 mM dNTPs (Life Technologies), 5 µM OligodT23VN (IDT), 10 µM DTT (Life Technologies) and 0.2 Units RNase inhibitor (SUPERase•In™, Invitrogen). RNA was denatured for 5 min at 65° C, before adding the reaction mix. Retrotranscription was performed at 42° C for 90 min, followed by 20 min at 70° C for enzyme inactivation. Standard RT-PCRs were run for all samples, including non-template controls, to corroborate amplification of single bands of the appropriate size. Cycling conditions were: 95° C for 3 min, (95 C, 15 s; 65-55° C (−0.5° C per cycle), 20 s; 68° C, 25 s), 20 cycles, (95° C, 15 s; 55° C, 20 s; 68° C, 25 s), 20 cycles, 68° C, 5 min, 4° C, hold. Reactions were run on a SimpliAmp™ Thermal Cycler (Applied Biosystems). Primers used were: ActB_For, cctctatgccaacacagtgc; ActB_Rev, cttctgcatcctgtcagcaa; Th_For, tgcctcctcacctatgcact; Th_Rev, gtcagccaacatgggtacg; Gad1_For, caagttctggctgatgtgga; Gad1_Rev, cttggcgtagaggtaatcagc; Tbp_For, agggattcaggaagaccacat; Tbp_Rev, cagtggtaccaaaactgactgg; cFos_For, agggagctgacagatacactcc; cFos_Rev, tgcaacgcagacttctcatc; Ddc_For, cacagaagtcattcttgggttg; Ddc_Rev, gagtttcgttcaactcattgga. Primers were designed using the Universal Probe Library Assay Design Centre (Roche), compatible with UPLs 157 (*ActB*), 3 (*Th*), 1 (*Gad1*), 2 (*Tbp*), 94 (*cFos*) and 38 (*Ddc*) primer pairs.

qPCR reactions were run using the FastStart Essential DNA Probes Master mix (Roche) on the LightCycler ® 96 Real-Time PCR System (Roche). Final primer concentration was 0.5 µM and probe concentration was 0.1 µM. Cycling conditions were: 95° C for 10 min, (95° C, 10 s; 60° C, 30 s), 55 cycles, 4° C, hold. Technical duplicates were performed for all samples and reactions. ActB was used as housekeeping reference gene for Th and Gad1, and Tbp for Ddc and cFos expression quantification. Amplification efficiency for all primer pairs was determined by obtaining a standard curve. Briefly, the cDNAs from each cDNA synthesis reaction were pooled and serially diluted. qPCR reactions were performed as above. Ct amplification curve values were plotted as a function of dilution factor and the amplification efficiency determined from the slope of the linear regression as in E = 10^(−1/slope)^-1. Amplification efficiency (E ± SE) for ActB primers was 1.07 ± 0.02, for Th primers, 1.01 ± 0.03, for Gad1 primers, 0.98 ± 0.04, for Tbp primers, 1.02 ± 0.02, for Ddc primers, 0.97 ± 0.04 and for cFos primers, 0.97 ± 0.04. Fold changes in Th, Gad1, cFos and Ddc expression levels were obtained using the 2^(–ddCt)^ method (Livak & Schmittgen, 2001; Taylor *et al*., 2019), first normalising Ct levels for each sample to housekeeping Ct levels and subsequently normalising to the average of the control (sham) conditions.

### Statistical analysis

Statistical analysis was carried out using Prism (GraphPad, San Diego, USA), Matlab (Mathworks, Massachusetts, USA) or SPSS (IBM). All datasets were described as mean ± SEM, unless otherwise stated, n values reported in the text refer to number of cells, and N values refer to number of mice. Details of all individual statistical tests are reported alongside the appropriate results. α values were set to 0.05 (except where a Bonferroni correction was carried out for multiple comparisons) and all comparisons were two tailed. For multiple group comparisons with multilevel ANOVA followed by Bonferroni post-tests, all groups either passed the D’Agostino and Pearson omnibus test for normality (GraphPad Prism), and/or could be fitted by a Gaussian distribution that explained ≥ 90 % of their variance (custom Matlab script written by MSG). Fisher’s exact test was performed when analysing proportions. Due to the design of this test, data had to be analysed twice, by combining groups based on manipulation and then duration of manipulation. A Bonferroni correction was therefore carried out for multiple comparisons in this instance (α = 0.05/2 = 0.025). qPCR normalised expression data were obtained from entirely separate 1 d (occluded versus control) and 3 d (occluded versus control) experiments, performed independently with different sets of reagents and with independent normalisation. For this reason we analysed these data with separate unpaired t-tests (with Welch’s correction where necessary where SDs were significantly different) for 1 day and 3 day samples. This therefore also required Bonferroni correction for multiple comparisons: again, α = 0.05/2 = 0.025.

## Results

### Using DAT-tdTomato mice to label olfactory bulb dopaminergic neurons

We used a conditional genetic strategy to label dopaminergic neurons in the olfactory bulb by crossing DAT^IRES*cre*^ mice (Bäckman *et al*., 2006) with a floxed tdTomato reporter line. This resulted in strong tdTomato fluorescence in small cells surrounding glomerular neuropil in the glomerular layer, along with the odd scattered tdTomato-positive cell in the superficial external plexiform layer (EPL), in keeping with previous reports of dopaminergic cell localisation within the olfactory bulb (Fig. 1A; (e.g. Hökfelt *et al*., 1975; Banerjee *et al*., 2013; Kosaka *et al*., 2019). To confirm the identity of these DAT-tdTomato neurons, we co-stained fixed olfactory bulb slices from juvenile postnatal day (P)28 mice for the rate-limiting enzyme in the synthesis of dopamine, tyrosine hydroxylase (TH). Although necessary for both dopamine and noradrenaline production, the absence of local noradrenergic cells in the olfactory bulb (Hökfelt *et al*., 1975; Rosser *et al*., 1986) makes TH a selective marker for dopaminergic neurons in this region. Previous analysis of this transgenic line found that ∼90 % of TH-positive glomerular layer neurons were also positive for tdTomato (Galliano et al., 2018); however, the composition of the tdTomato-positive population has not yet been fully described. In control animals we found a high incidence of co-labelled neurons, with ∼75% of DAT-tdTomato cells staining positively for TH (Fig. 1B,C; % of tdTomato-positive neurons ± SEM: TH-positive, 74 ± 4 %; TH-negative, 26 ± 4 %; n = 598 cells, N = 3; Vaaga *et al*., 2017). However, by no means all genetically labelled cells in the DAT-tdTomato mouse were TH-positive. We reasoned that the ∼25 % of non-TH-expressing DAT-tdTomato cells might be ‘true’ dopaminergic neurons that have low, undetectable levels of TH expression due to long-term activity-dependent down-regulation (e.g. Baker *et al*., 1983). However, co-staining for another dopamine-synthesising enzyme dopa decarboxylase (DDC), the expression of which is known to be resistant to long-term changes in ongoing activity (Stone *et al*., 1990; Baker *et al*., 1993; Cave *et al*., 2010), showed that this is not the case. We found very good agreement between TH- and DDC-positive cells in the olfactory bulb glomerular layer (Fig. 1B; 99 ± 0.5 % of TH-positive neurons were also DDC-positive; 89 ± 2 % of DDC-positive neurons were also TH-positive; n = 598; N = 3), but we found only 3 % of DAT-tdTomato cells were positive for DDC without expressing TH (Fig. 1B,C; % of tdTomato-positive neurons ± SEM: TH- and DDC-positive, 74 ± 4 %; DDC-positive but TH-negative, 3 ± 7 %; TH-positive but DDC-negative, 0.5 ± 0.3 %; TH- and DDC-negative, 23 ± 2 %; n = 598; N = 3).

If ∼20-25 % of DAT-tdTomato cells in the olfactory bulb are therefore non-dopaminergic, what type of cell are they? Red fluorescent neurons never had the large, pear-shaped soma characteristic of local excitatory neurons, the external tufted cells, in the glomerular layer (Hayar *et al*., 2004). This made them most likely to be GABAergic juxtaglomerular interneurons, which in mouse are divided into three mutually exclusive subgroups expressing either TH or the calcium-binding proteins calbindin and calretinin (Kohwi *et al*., 2007; Kosaka & Kosaka, 2007; Panzanelli *et al*., 2007; Parrish-Aungst *et al*., 2007). We never observed any DAT-tdTomato neurons that co-labelled for calbindin (Fig. 1D), but we did find DAT-tdTomato cells that were positive for calretinin (Fig. 1E; Ninkovic *et al*., 2010). Indeed, the tdT-positive/TH-negative subgroup was almost entirely accounted for by cells expressing calretinin (Fig. 1E, F; % of tdTomato-positive neurons ± SEM: TH-positive only, 74 ± 2 %; calretinin-positive only, 22 ± 0.9 %; n = 1404; N = 3). Importantly, although we observed some rare tdTomato-positive cells that were neither TH-nor calretinin-positive (Fig. 1F; tdTomato-positive only, 4 ± 1%, consistent with the 3 % of tdTomato cells found above to be DDC-positive but TH-negative), we never saw any tdTomato-positive cells co-labelled for both TH and calretinin (n = 0 / 1404 cells from N = 3 mice). This suggests that the calretinin-positive DAT-tdTomato cells are non-dopaminergic calretinin neurons that have at least transiently activated DAT expression at some point in their history, rather than being dopaminergic neurons that also co-express calretinin.

To check these cells’ distinct properties, we obtained whole-cell patch-clamp recordings of DAT-tdTomato neurons in acute olfactory bulb slices. Although these cells are structurally fragile and extremely difficult to recover for post-recording histology (Galliano *et al*., 2018), we were able to find very rare example neurons that had been successfully filled with biocytin during our recordings and then co-labelled for glomerular layer markers. While a confirmed TH-positive DAT-tdTomato neuron had characteristic multiple spike production in response to a prolonged 500 ms current injection in current-clamp, the one calretinin-positive tdTomato cell we managed to recover displayed only a single, small spike followed by a sustained depolarised plateau potential, as previously described to be typical of this bulbar cell type (Fig. 1G; Fogli Iseppe *et al*., 2016; Benito *et al*., 2018). The calretinin-positive cells labelled in DAT-tdTomato mice are therefore likely to share their major features with the general population of calretinin-expressing juxtaglomerular neurons.

### Sensory deprivation via brief unilateral naris occlusion reduces cFos expression in olfactory bulb DAT-tdTomato neurons

Having shown that the DAT-tdTomato mouse line labels olfactory bulb neurons that are predominantly dopaminergic but include a distinct subset of calretinin-positive cells, we went on to use this line for naturally-relevant manipulations of sensory experience (Fokkens *et al*., 2012; Galliano et al., 2021). In juvenile mice from postnatal day (P)27, we induced brief 1- or 3-day periods of sensory deprivation without damaging the olfactory epithelium, via unilateral naris occlusion with a plastic plug (Fig.2A; Cummings *et al*., 1997; Kikuta *et al*., 2015; Cheetham *et al*., 2016; Galliano et al., 2021). Because of interpretational issues caused by abnormal airflow through the contralateral naris in such manipulations (Coppola, 2012; Kass *et al*., 2013), we did not compare deprived versus spared olfactory bulbs in the same animals, and instead made all comparisons between the right bulbs of individual occluded or control mice (see Methods).

**Figure 2.**
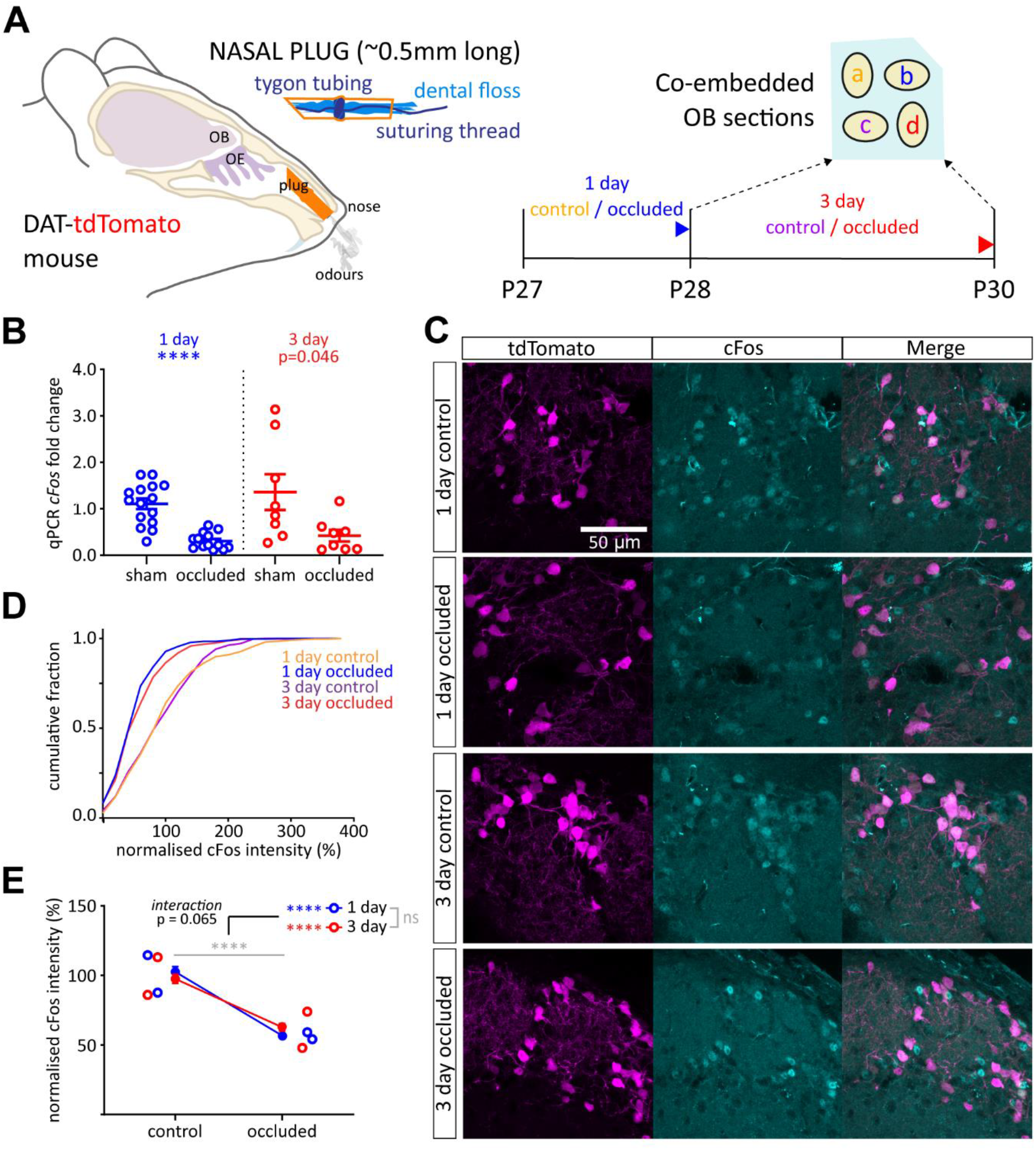
Sensory deprivation via brief unilateral naris occlusion reduces cFos expression in olfactory bulb DAT-tdTomato neurons. **A)** Schematic of naris occlusion protocol and tissue co-embedding for comparison of relative immunofluorescence. OE, Olfactory epithelium; OB, Olfactory bulb; P, Postnatal day; a, 1 day control; b, 1 day occluded; c, 3 day control; d, 3 day occluded. **B)** *cFos* mRNA expression by whole-bulb qPCR. Circles show fold change for individual bulbs vs relevant sham control mean; lines show mean ± SEM fold change for each group. t-test (α = 0.025); 1 day, ****, p < 0.0001; 3 day, p = 0.046. **C)** Example maximum intensity projection images showing immunofluorescent label for cFos in the glomerular layer of the olfactory bulb in 1 and 3 day control and occluded DAT-tdTomato mice. All images are from the same co-embedded preparation, acquired and presented with identical settings. **D)** Cumulative distribution of normalised cFos intensity (%) in DAT-tdTomato neurons in each treatment group. **E)** Line plots showing overall mean ± SEM normalised cFos intensity (%) in each treatment group; unfilled circles show mean normalised cFos intensity (%) for individual mice. Multi-level ANOVA controlling for set variance; effect of manipulation, **** p < 0.0001; effect of time, ns: non-significant. Bonferroni post-test (α = 0.025); 1 day, effect of manipulation, **** p < 0.00005; 3 day, effect of manipulation, **** p < 0.00005

To first validate the effectiveness of our sensory deprivation approach for reducing overall olfactory bulb activity, we assessed levels of whole-bulb mRNA for the immediate early gene *cFos* using qPCR in wild-type mice. We found decreases in *cFos* mRNA after both 1 day or 3 day naris occlusion compared to time-matched sham controls (Fig.2B). The 1 day effect was highly significant, while variability within the 3 day control group meant that the effect of occlusion at this time point fell just short of significance after correction for multiple comparisons (1 day mean ± SEM fold change: sham, 1.11 ± 0.12; occluded, 0.31 ± 0.046; Welch’s t-test with α = 0.025, t_18.25_ = 6.45, p < 0.0001, N = 29; 3 day: sham, 1.36 ± 0.38; occluded, 0.42 ± 0.13; Welch’s t-test with α = 0.025, t_8.50_ = 2.33, p = 0.046; N = 16).

To specifically assess the effect of brief sensory deprivation on spontaneous activity levels in DAT-tdTomato neurons, we then examined cFos immunofluorescence (Liu *et al*., 1999). Co-embedding olfactory bulbs from all four treatment groups (Fig. 2A; 1 day control, 1 day occluded; 3 day control; 3 day occluded) prior to sectioning allowed the best possible consistency in immunostaining protocols (see Methods; Vlug *et al*., 2005; Galliano et al., 2021). It also enabled us to normalise labelling intensity within co-embedded sets in order to combine analyses across multiple individuals. With this approach we found that DAT-tdTomato cells from occluded mice had significantly lower cFos immunofluorescence after both 1 and 3 days of occlusion (Fig. 2C-E; normalised intensity (%) mean ± SEM: 1 day control, 102.50 ± 3.83%, n = 316 cells, N = 2 mice; 1 day occluded, 56.85 ± 2.10%, n = 358, N = 2; 3 day control, 97.53 ± 3.30%, n = 317, N = 2; 3 day occluded, 62.87 ± 2.63%, n = 309, N = 2. Multi-level ANOVA controlling for set variance, effect of manipulation, F_1,1298_ = 60.73, p < 0.0001; effect of time, F_1,1298_ = 1.40, p = 0.24; effect of manipulation × time interaction, F_1,1299_ = 3.40, p = 0.065. Bonferroni post-test comparing control versus occluded (α = 0.025): 1 day, F_1,673_ = 115.76, p < 0.00005; 3 day, F_1,626_ = 68.56, p < 0.00005). Brief sensory deprivation therefore reliably lowers activity levels in olfactory bulb DAT-tdTomato neurons, and does so in a sustained manner over 1 to 3 days.

### Brief naris occlusion does not alter the sub-type composition of olfactory bulb DAT-tdTomato neurons

Given the mixed dopaminergic and calretinin-expressing nature of the population labelled in the DAT-tdTomato mouse olfactory bulb (Fig. 1), it was important to check whether our effective short-term sensory manipulations (Fig. 2) altered this sub-type composition in any way. Crucially, we found no change in the proportion of DAT-tdTomato cells that were calretinin-positive after 1 or 3 days of sensory deprivation (Fig. 3A,B; % of calretinin-positive DAT-tdTomato neurons; 1 day control, 20.95 %, n = 506, N = 3; 1 day occluded, 24.50 %, n = 502, N = 3; 3 day control, 23.91 %, n = 343, N = 2; 3 day occluded, 23.74 %, n = 417, N = 3. Fisher’s exact test with α = 0.025, effect of manipulation, p = 0.22; effect of time, p = 0.40).

**Figure 3.**
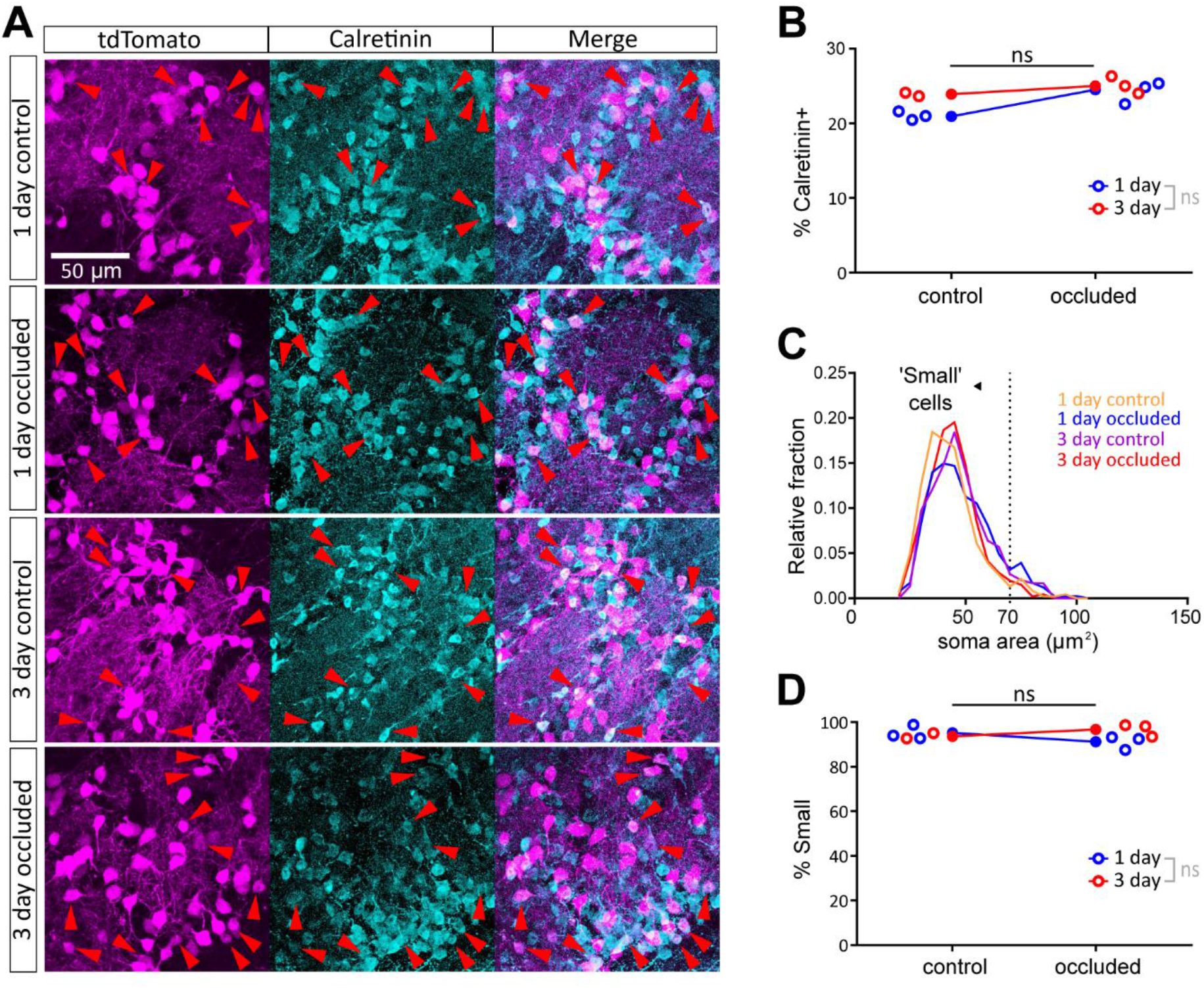
Brief naris occlusion does not alter the sub-type composition of olfactory bulb DAT-tdTomato neurons. **A)** Example maximum intensity projection images showing immunofluorescence for tdTomato and calretinin in the glomerular layer of the olfactory bulb in 1 and 3 day control and occluded DAT-tdTomato mice. Red arrowheads indicate example neurons co-expressing DAT-tdTomato and calretinin. **B)** Line plots showing, for each treatment group, overall % of tdTomato+ neurons that also express calretinin. Open circles show % values for individual mice. Fisher’s exact test with Bonferroni correction for multiple comparisons (α = 0.025); ns, non-significant. **C)** Histogram of soma sizes for all tdTomato+ neurons in each treatment condition. Dotted line shows 70 µm^2^ threshold for cells classed as ‘small’. **D)** Small cell % data; all conventions as in **B**.

Olfactory bulb dopaminergic neurons are not a unitary, homogeneous cell type (Galliano *et al*., 2018; Kosaka *et al*., 2019), so it was also important to assess the representation of dopaminergic subtypes amongst DAT-tdTomato neurons following sensory deprivation. Work from our laboratory and others has identified a key distinction between, on the one hand, small-soma olfactory bulb dopaminergic cells that lack an axon and can be generated throughout life and, on the other, large-soma olfactory bulb dopaminergic cells that possess an axon and are exclusively generated in early embryonic development (Galliano *et al*., 2018; Kosaka *et al*., 2019). The latter large-soma sub-population, although present, is significantly under-represented by DAT-tdTomato label, which is dominated by small-soma anaxonic cells (Galliano *et al*., 2018). Using a previously established soma area cut-off value of 70 µm^2^ to define olfactory bulb dopaminergic cells as ‘small’ (Galliano *et al*., 2018), we confirmed here that this subtype accounts for > 90 % of DAT-tdTomato neurons (Fig.3C,D). Crucially, the high proportion of such small cells represented in the DAT-tdTomato line – which will also include the small-soma calretinin-positive subgroup (Parrish-Aungst *et al*., 2007) – was not significantly affected by 1 or 3 day naris occlusion (Fig.3D; % of ‘small’ cells; 1 day control, 95.22 %, n = 460, N = 3; 1 day occluded, 91.18 %, n = 408, N = 3; 3 day control, 93.62 %, n = 298, N = 2; 3 day occluded, 96.75 %, n = 461, N = 3. Fisher’s exact test with α = 0.025, effect of manipulation, p = 0.75; effect of time, p = 0.067). Brief periods of sensory deprivation therefore do not significantly alter the cellular sub-composition of olfactory bulb glomerular layer neurons labelled in DAT-tdTomato mice.

### Brief sensory deprivation does not affect GAD67 expression in olfactory bulb DAT-tdTomato neurons

Having confirmed that brief unilateral naris occlusion effectively lowers activity levels in the olfactory bulb DAT-tdTomato population without affecting its subtype composition, we used this mouse line to investigate potential experience-dependent plasticity in the expression of neurotransmitter-synthesising enzymes. We focused first on the GABA-synthesising enzyme isoform GAD67, which is ubiquitously expressed in dual GABA- and dopamine-releasing olfactory bulb neurons (Kosaka & Kosaka, 2007; Kiyokage *et al*., 2010; Kosaka *et al*., 2019), and has been previously reported to undergo experience-dependent changes in both mRNA expression and protein levels after prolonged manipulations (Parrish-Aungst *et al*., 2011; Lau & Murthy, 2012; Banerjee *et al*., 2013; Wang *et al*., 2017). However, here we observed no effect of either 1 day or 3 day naris occlusion on levels of *Gad1* mRNA from wild-type whole-bulb samples (Fig.4A; 1 day mean ± SEM fold change: sham, 1.06 ± 0.12; occluded, 1.08 ± 0.082; t-test with α = 0.025, t_18_ = 0.11, p = 0.92, N = 20; 3 day: sham, 1.01 ± 0.041; occluded, 0.90 ± 0.061; t-test with α = 0.025, t_14_ = 1.45, p = 0.17; N = 16). We also found no changes in relative GAD67 immunofluorescence intensity in olfactory bulb DAT-tdTomato neurons after 1 or 3 days of unilateral naris occlusion (Fig.4B-D; normalised immunofluorescence intensity (%) mean ± SEM: 1 day control, 97.43 ± 3.08 %, n = 261, N = 3; 1 day occluded, 92.93 ± 2.89 %, n = 314, N = 3; 3 day control, 103.50 ± 3.70 %, n = 192, N = 2; 3 day occluded, 94.03 ± 2.88 %, n = 304, N = 3. Multi-level ANOVA controlling for set variance, effect of manipulation, F_1,1071_ = 0.089, p = 0.77; effect of time, F_1,1071_ = 1.60, p = 0.21; effect of manipulation × time interaction, F_1,1071_ = 0.62, p = 0.43).

**Figure 4.**
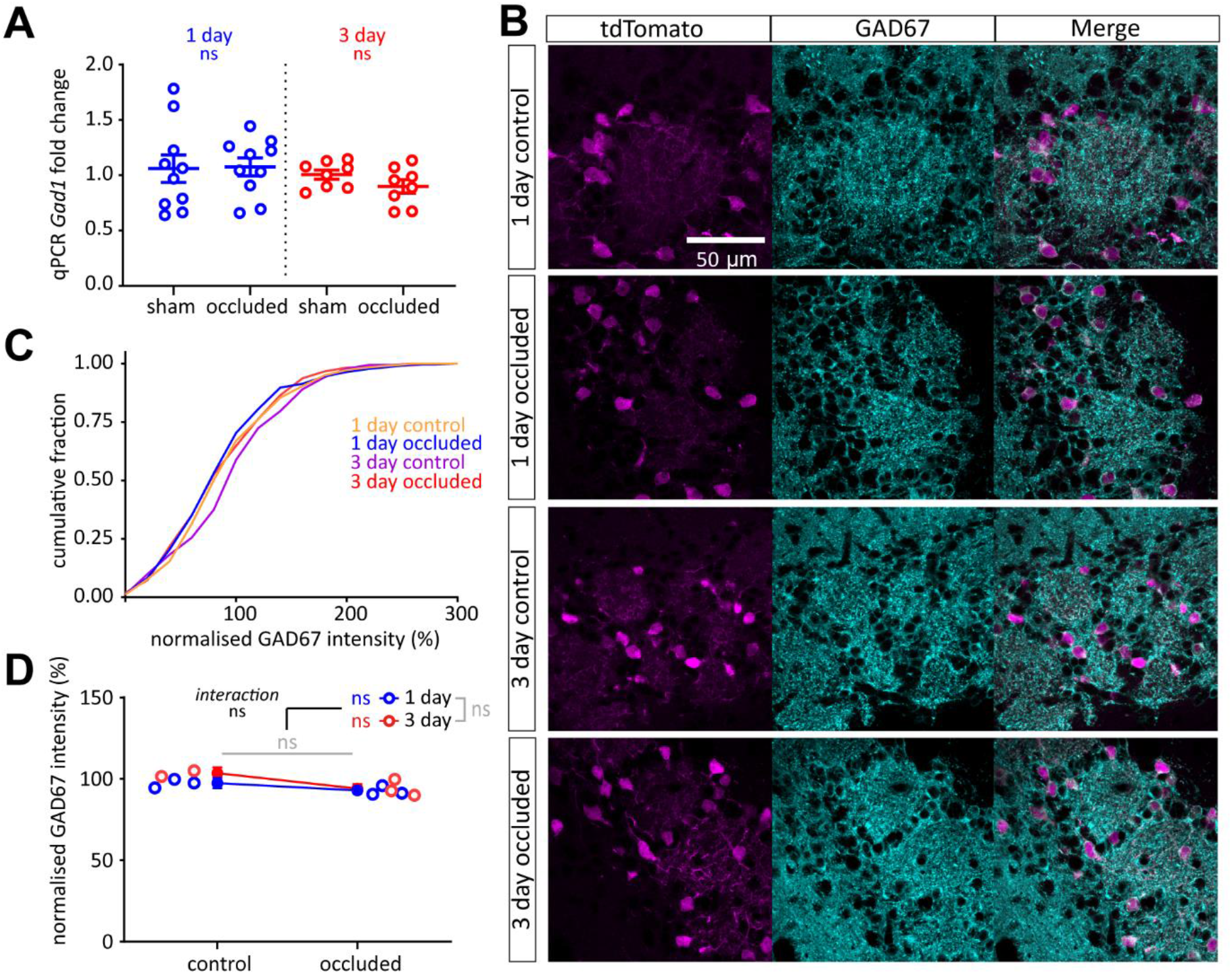
Brief sensory deprivation does not affect GAD67 expression in olfactory bulb DAT-tdTomato neurons. **A)** *Gad1* mRNA expression by whole-bulb qPCR. Circles show fold change for individual bulbs vs relevant sham control mean; lines show mean ± SEM fold change for each group. **B)** Example maximum intensity projection images showing immunofluorescence for GAD67 in the glomerular layer of the olfactory bulb in 1 and 3 day control and occluded DAT-tdTomato mice. All images are from the same co-embedded preparation, acquired and presented with identical settings. **(C)** Cumulative distribution of normalised GAD67 intensity (%) in DAT-tdTomato neurons in each treatment group. **D)** Line plots showing overall mean ± SEM of normalised GAD67 intensity (%) in each treatment group; unfilled circles show mean normalised GAD67 intensity (%) for individual mice. Multi-level ANOVA controlling for set variance; effect of manipulation and effect of time, ns: non-significant.

### DDC expression in olfactory bulb DAT-tdTomato neurons is only slightly and transiently reduced by unilateral naris occlusion

The enzyme DDC catalyses the direct synthesis of dopamine from L-DOPA, but unlike TH has received relatively little attention regarding its potential for activity-dependent expression. Indeed, two previous studies found no effect of odour deprivation on *Ddc* mRNA levels in the olfactory bulb following prolonged, >1 month-long naris closure (Stone *et al*., 1990; Cave *et al*., 2010). Here, in whole-bulb samples from wild-type mice, we observed a trend towards increased *Ddc* mRNA levels after 1 day naris occlusion, but no effect of sensory deprivation at the longer, 3 day interval (Fig.5A; 1 day mean ± SEM fold change: sham, 1.08 ± 0.11; occluded, 1.35 ± 0.099; t-test with α = 0.025, t_27_ = 1.81, p = 0.081, N = 29; 3 day: sham, 1.03 ± 0.11; occluded, 1.14 ± 0.16; t-test with α = 0.025, t_14_ = 0.52, p = 0.61; N = 16). In contrast, examining relative DDC immunofluorescence intensity specifically in olfactory bulb DAT-tdTomato neurons revealed a very small but nevertheless significant *decrease* after naris occlusion. (Fig. 5B-D; normalised immunofluorescence intensity (%) mean ± SEM: 1 day control, 99.65 ± 1.44%, n = 884, N = 4; 1 day occluded, 92.17 ± 1.51%, n = 841, N = 4; 3 day control, 100.51 ± 1.48%, n = 609, N = 3; 3 day occluded, 98.56 ± 1.53%, n = 819, N = 4. Multi-level ANOVA controlling for set variance, effect of manipulation, F_1,3152_ = 10.62, p = 0.0010; effect of time, F_1,3097_ = 0.00, p = 0.99; effect of manipulation × time interaction, F_1,3140_ = 4.75, p = 0.029). The significant manipulation-by-time interaction here was supported by pairwise post-hoc tests demonstrating that this effect was driven by a significant reduction in DDC immunofluorescence only after 1 day of occlusion, with expression of DDC in 3 day occluded DAT-tdTomato neurons then returning back to control levels (Fig. 5D; Bonferroni post-test comparing control versus occluded with α = 0.025: 1 day, F_1,1722_ = 11.84, p < 0.0001; 3 day, F_1,738_ = 0.59, p = 0.44). Small and transient changes in relative DDC levels are therefore possible in olfactory bulb dopaminergic cells experiencing sustained decreases in activity.

**Figure 5.**
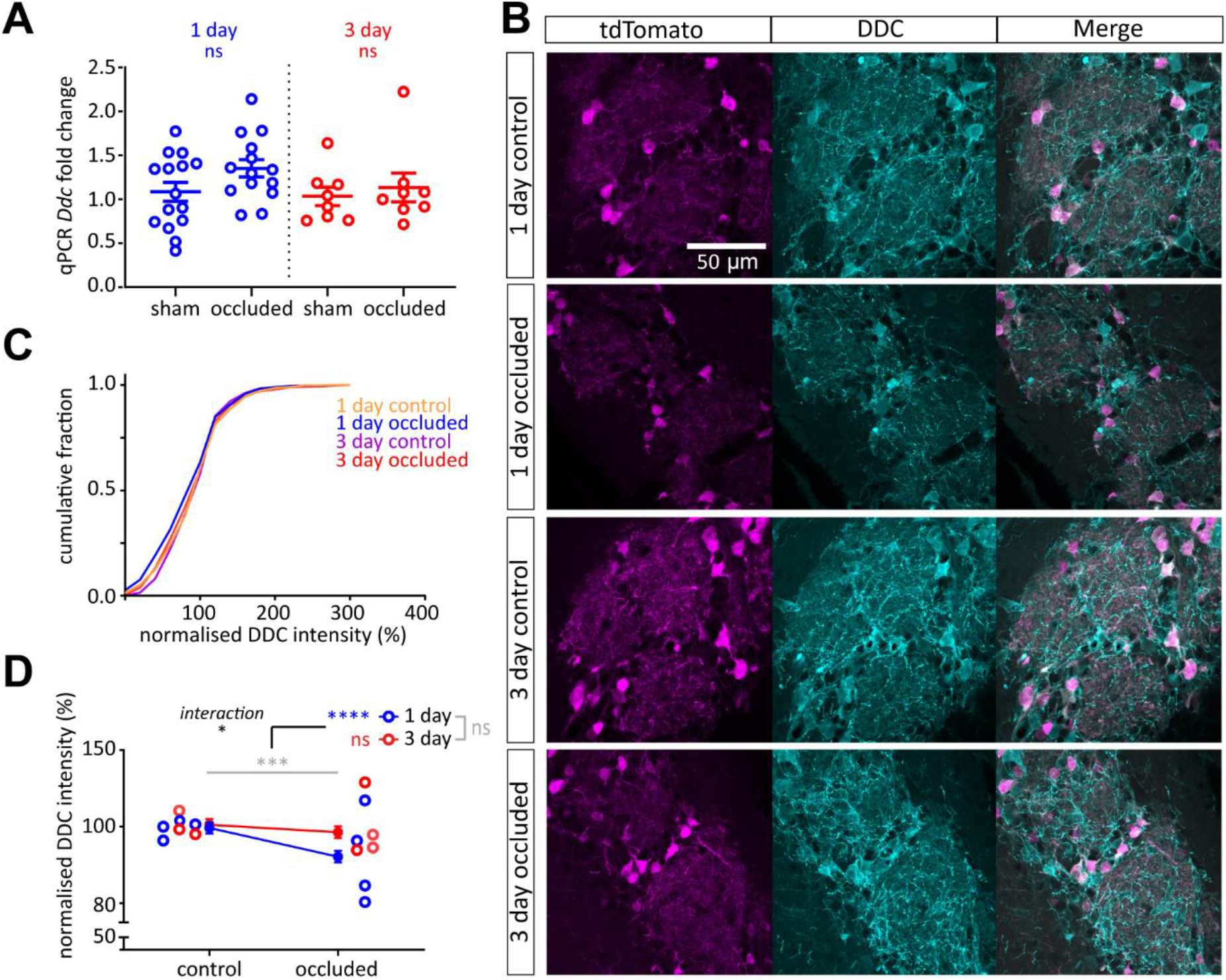
DDC expression in olfactory bulb DAT-tdTomato neurons is slightly reduced only after 1 day of unilateral naris occlusion. **A)** *Ddc* mRNA expression by whole-bulb qPCR. Circles show fold change for individual bulbs vs relevant sham control mean; lines show mean ± SEM fold change for each group. **B)** Example maximum intensity projection images showing immunofluorescence for DDC in the glomerular layer of the olfactory bulb in 1 and 3 day control and occluded DAT-tdTomato mice. All images are from the same co-embedded preparation, acquired and presented with identical settings. **C)** Cumulative distribution of normalised DDC intensity (%) in DAT-tdTomato neurons in each treatment group. **(D)** Line plots showing overall mean ± SEM of normalised DDC intensity (%) in each treatment group; unfilled circles show mean normalised DDC intensity (%) for individual mice. Multi-level ANOVA controlling for set variance; effect of manipulation, *** p < 0.001; effect of time, ns: non-significant; effect of interaction, * p < 0.05. Bonferroni post-test (α = 0.025); 1 day, effect of manipulation, **** p < 0.0001; 3 day, effect of manipulation, ns: non-significant.

### Brief sensory deprivation reduces TH expression in olfactory bulb DAT-tdTomato neurons

Olfactory bulb dopaminergic neurons have a well-described ability to regulate TH expression in an activity-dependent manner, including after unilateral naris occlusion by plug insertion (Nadi *et al*., 1981; Baker *et al*., 1983; Kosaka *et al*., 1987; Cummings *et al*., 1997). At the mRNA level, plasticity in *Th* expression has been previously reported after 2 days of odour deprivation (Cho *et al*., 1996). Here we also observed rapid occlusion-induced downregulation of *Th* mRNA levels, as assessed via qPCR on whole-bulb samples from wild-type mice. Compared to sham-treated bulbs, OBs on the occluded side had significantly lower levels of *Th* expression after both 1 day and 3 days of sensory deprivation (Fig.6A; 1 day mean ± SEM fold change: sham, 1.02 ± 0.049; occluded, 0.64 ± 0.030; t-test with α = 0.025, t_27_ = 6.55, p < 0.0001, N = 29; 3 day: sham, 1.04 ± 0.13; occluded, 0.53 ± 0.029; Welch’s t-test with α = 0.025, t_7.73_ = 4.00, p = 0.0043; N = 16).

**Figure 6.**
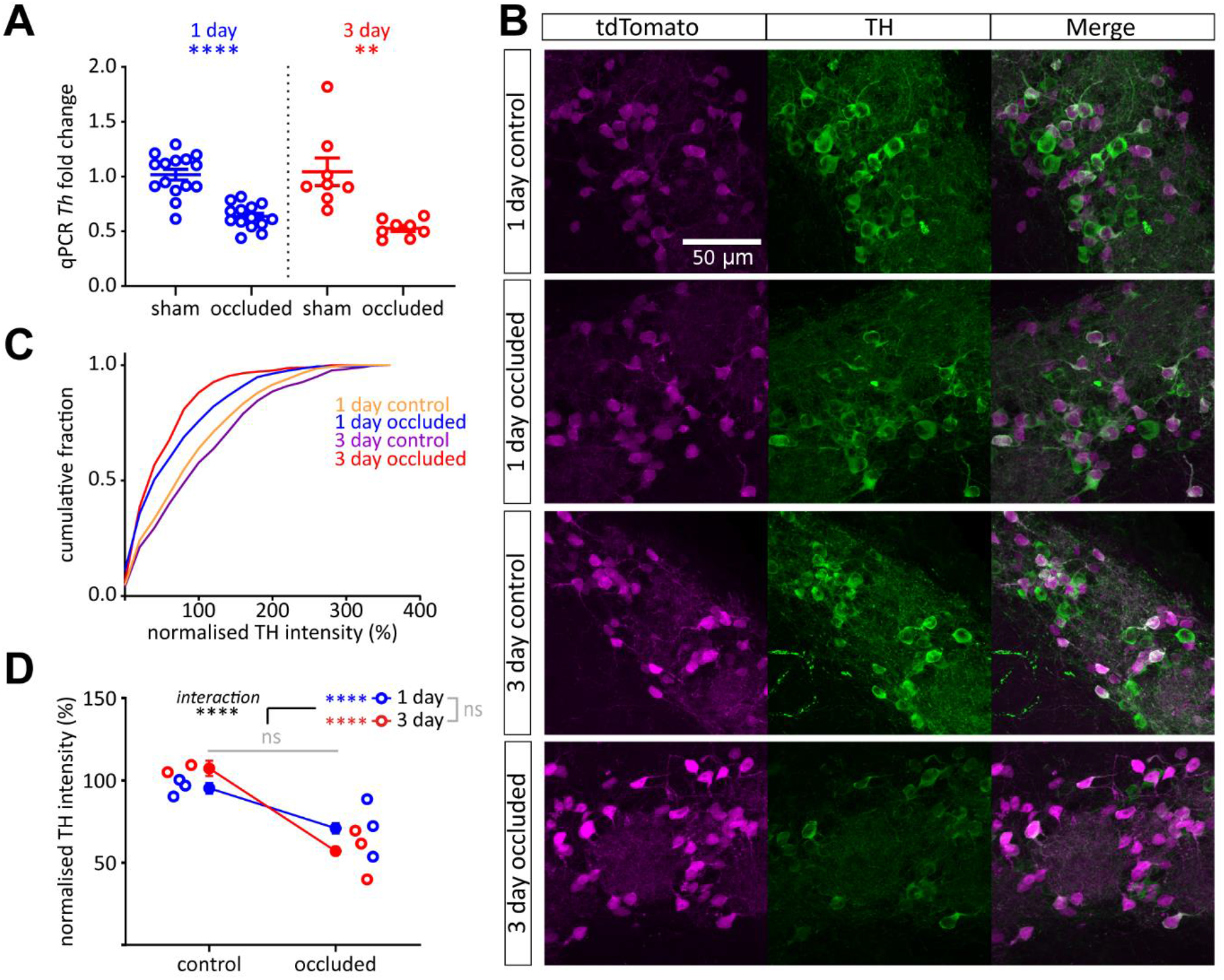
Brief sensory deprivation reduces TH expression in olfactory bulb DAT-tdTomato neurons. **A)** *Th* mRNA expression by whole-bulb qPCR. Circles show fold change for individual bulbs vs relevant sham control mean; lines show mean ± SEM fold change for each group. t-test (α = 0.025); 1 day, ****, p < 0.0001; 3 day, **, p = 0.0043. **B)** Example maximum intensity projection images showing immunofluorescence for TH in the glomerular layer of the olfactory bulb in 1 and 3 day control and occluded DAT-tdTomato mice. All images are from the same co-embedded preparation, acquired and presented with identical settings. **C)** Cumulative distribution of normalised TH intensity (%) in DAT-tdTomato neurons in each treatment group. **D)** Line plots showing overall mean ± SEM of normalised TH intensity (%) in each treatment group; unfilled circles show mean normalised TH intensity (%) for individual mice. Multi-level ANOVA controlling for set variance; effect of manipulation × time interaction, **** p < 0.0001; effect of manipulation and effect of time, ns: non-significant. Bonferroni post-test (α = 0.025); 1 day, effect of manipulation, **** p < 0.00005; 3 day, effect of manipulation, **** p < 0.00005.

At the protein level, just one day of elevated activity in cultured bulbar neurons or slices has been found to increase TH immunofluorescence intensity or TH-GFP transgene expression (Akiba *et al*., 2007; Chand *et al*., 2015). We also recently found that 24 h naris occlusion was accompanied by a significant drop in immunofluorescence for TH in the ipsilateral olfactory bulb, without producing a change in TH-positive cell density (Galliano et al., 2021). However, the above approaches had the drawback that they used TH expression itself as a means to identify samples for TH quantification. We therefore used our conditional genetic labelling strategy to assess rapid deprivation-induced changes in TH expression in an independently identified neuronal population.

Brief sensory deprivation was indeed associated with a decrease in the relative intensity of TH immunofluorescence in olfactory bulb DAT tdTomato neurons (Fig.6B-D; normalised immunofluorescence intensity (%) mean ± SEM: 1 day control, 95.27 ± 3.35%, n = 460, N = 3; 1 day occluded, 70.96 ± 3.16%, n = 408, N = 3; 3 day control, 107.30 ± 4.64%, n = 298, N = 2; 3 day occluded, 57.00 ± 2.32%, n = 461, N = 3. Multi-level ANOVA controlling for set variance, effect of manipulation, F_1,1546._ = 2.90, p = 0.089; effect of time, F_1,1626_ = 3.10, p = 0.078). This effect was highly significant after just 1 day (Fig. 6C; Bonferroni post-test comparing control versus occluded with α = 0.025: 1 day, F_1,867_ = 27.41, p < 0.00005), but was even stronger after 3 days (Fig. 6C; post-test: 3 day, F_1,639_ = 76.36, p < 0.00005; effect of manipulation × time interaction in main ANOVA, F_1,1613_ = 11.67, p = 0.001). Continuous naris occlusion therefore produces a rapid downregulation in TH immunofluorescence in independently identified olfactory bulb dopaminergic neurons, and this effect becomes stronger over time.

## Discussion

Here we used a conditional genetic strategy to identify olfactory bulb neurons expressing the dopamine transporter. DAT-tdTomato neurons were predominantly dopaminergic, but also included a subset of non-dopaminergic, calretinin-positive cells. Importantly, while cFos immunolabel showed that the activity of DAT-tdTomato cells could be readily manipulated with brief periods of sensory deprivation, these manipulations did not alter the labelled neurons’ subtype composition. We found that 1-3 days of unilateral naris occlusion was insufficient to alter expression of the GABA-synthesising enzyme GAD67 in these cells. In contrast, similarly brief sensory manipulations were associated with significant activity-dependent changes in the levels of enzymes required for the synthesis of dopamine.

### Identifying olfactory bulb dopaminergic neurons using DAT^IREScre^ mice

Using transgenic approaches to label specific cell types in the brain requires thorough characterisation of their specificity. To specifically label dopaminergic neurons in the olfactory bulb’s glomerular layer, several previous studies have successfully utilised conditional or knock-in strategies based on expression driven from the *Th* promoter (e.g. Saino-Saito *et al*., 2004; Pignatelli *et al*., 2005; Kiyokage *et al*., 2010; Adam & Mizrahi, 2011; Liu *et al*., 2013; Wachowiak *et al*., 2013). However, this approach can also result in labelled presumptive immature migrating neurons outside the glomerular layer, which can express *Th* mRNA before TH protein expression is detectable (Saino-Saito *et al*., 2004; Pignatelli *et al*., 2009; Banerjee *et al*., 2013). We therefore opted for a different conditional genetic labelling strategy based on Cre recombinase expression under the control of the DAT promoter, which has been reported to be specific to TH-positive neurons in general (Lammel *et al*., 2015), and which has already been used successfully to target olfactory bulb dopaminergic neurons. In those bulb studies, which either employed the DAT^IRES*cre*^ line used here (Vaaga *et al*., 2017; Short & Wachowiak, 2019; Sato *et al*., 2020) a DAT-Cre knock-in model (Ninkovic *et al*., 2010; Banerjee *et al*., 2015), or a BAC transgenic DAT-CreER^T2^ inducible line (Bywalez *et al*., 2016), approximately 85% of genetically-labelled neurons were found to be TH-positive by immunohistochemistry (Banerjee *et al*., 2015; Vaaga *et al*., 2017). This is broadly in line with the ∼75% co-label we found here, with cross-study variability possibly arising from differences in the Cre-dependent reporter lines utilised and/or differences in label detection criteria. Nevertheless, the general approach of using DAT-Cre mice to label dopaminergic neurons in the olfactory bulb clearly produces reasonable specificity.

However, we found that not all DAT-tdTomato olfactory bulb cells were TH-positive, and that the large majority of non-dopaminergic genetically labelled neurons (∼20 %) expressed the calcium-binding protein calretinin. Calretinin expression in DAT-Cre-positive bulbar neurons has been previously reported, but not quantified, in a separate knock-in line (Ninkovic *et al*., 2010). This suggests that off-target label of calretinin cells is not a peculiarity of one particular genetic manipulation, but instead reflects a biological link between these neurons and their dopaminergic neighbours. Indeed, fate mapping neuronal precursors that generate glomerular layer GABAergic interneurons has demonstrated that the decision to differentiate into a dopaminergic neuron over a calretinin neuron relies on just a few transcription factors. Loss of Pax6, Dlx2 or Meis2 reduces the proportion of postnatal dopaminergic neurons that are generated, while increasing the number of calretinin neurons (Brill *et al*., 2008; Agoston *et al*., 2014). Overexpression of Dlx2 increases the proportion of dopaminergic neurons and decreases the calretinin population (Brill *et al*., 2008). The transcription factor Er81 is also expressed in both calretinin and dopaminergic neurons in the glomerular layer and is important in mice for regulating the expression of *Th* (Cave *et al*., 2010). Therefore, the molecular determinants of dopaminergic versus calretinin identities appear to be closely interdependent. It remains unclear whether tdTomato expression in some calretinin neurons is due to transient expression of DAT during their development, or whether this subset of cells continues to express DAT postnatally. In the former case, obtaining transgenic label via inducible activation of DAT-Cre in mature mice (Bywalez *et al*., 2016) might result in greater specificity for bulbar dopaminergic neurons. Low levels of bulbar DAT expression (Cockerham *et al*., 2016) unfortunately precluded us from addressing this question using immunohistochemistry, but targeted interrogation of single-cell sequencing datasets (e.g. Zeisel *et al*., 2018) might provide an answer in the near future. Regardless, it is clear that this minority calretinin-expressing population needs to be considered when interpreting the results of experiments performed on ‘dopaminergic’ neurons targeted via DAT-Cre mouse lines (Wachowiak *et al*., 2013; Banerjee *et al*., 2015; Bywalez *et al*., 2016; Vaaga *et al*., 2017; Galliano *et al*., 2018, 2020; Sato *et al*., 2020). Thankfully, though, the distinct and unusually immature-like properties of glomerular layer calretinin-positive cells, including their extremely low excitability and apparent lack of GABA release (Fig. 1G; Fogli Iseppe *et al*., 2016; Benito *et al*., 2018; Tavakoli *et al*., 2018), mean that their presence as a minority of DAT-Cre-targeted cells is unlikely to have significantly biased the conclusions of those previous studies.

Our main goal was to achieve reasonable specificity in genetically labelling olfactory bulb dopaminergic neurons, but we note that our approach was by no means comprehensive in labelling *all* olfactory bulb dopaminergic neurons. As is the case for many Cre lines (Heffner *et al*., 2012), we saw mosaicism in the fact that many immunohistochemically-defined TH-positive neurons did not express the tdTomato marker (see Fig. 1B,E; % of TH-positive neurons that were tdTomato-negative, 36 ± 0.2 %, n = 692 cells, N = 3 mice; Galliano *et al*., 2018). The DAT^IRES*cre*^ line is also known to particularly under-represent one subtype of bulbar dopaminergic neuron – the large-soma, axon-bearing, embryonically-generated subclass (Galliano *et al*., 2018) – and we found again here that small-soma, presumed anaxonic neurons dominate the tdT-labelled population. However, large-soma, axon-positive cells are already rare amongst the olfactory bulb dopaminergic population, and they can certainly still be labelled in DAT-tdTomato mice (Galliano *et al*., 2018). So, although the DAT^IRES*cre*^ line produces genetic label which is heavily biased towards one particular dopaminergic subpopulation in the olfactory bulb, it is far from being entirely subtype specific. Overall, the strategy of using DAT-based transgenic lines to target dopaminergic neurons in the olfactory bulb can produce good levels of specificity, but care needs to be taken to fully characterise the resulting label before such experiments can be safely interpreted.

### Experience-dependent plasticity of GAD67

With brief periods of sensory deprivation, we saw no evidence here for changes in relative GAD67 immunofluorescence amongst olfactory bulb DAT-tdTomato-expressing neurons. Given the mixed calretinin- and TH-positive composition of our tdTomato-labelled population (Fig. 1), this result could theoretically have been produced if changes in GAD67 levels in dopaminergic neurons were perfectly offset by inverse, and stronger, changes in the smaller calretinin-positive tdTomato subgroup. However, such effects might be expected to significantly change the shape of the distribution of GAD67 relative immunofluorescence values, and we saw no obvious changes of this nature in our data (Fig.4C). We did not co-label for calretinin and GAD67 in this study, while the literature contains conflicting reports as to whether calretinin-positive neurons in the olfactory bulb express GAD67 at all (Panzanelli *et al*., 2007; Parrish-Aungst *et al*., 2007; opposing results are likely due to the different transgenic versus immunohistochemical labelling methods employed in these studies). Even if calretinin-positive tdTomato cells do express GAD67, however, the most parsimonious interpretation of our data remains that dopaminergic bulbar neurons do not rapidly alter their GAD67 levels in response to sensory deprivation. This fits with previous work measuring general GAD (i.e. GAD65+67) expression in the bulb, which found no significant alterations following more long-term sensory deprivation (Kosaka *et al*., 1987; Stone *et al*., 1991; Baker *et al*., 1993). In contrast, reduced expression of GAD67, but not GAD65, has been observed after 7 days of unilateral naris occlusion and deafferentation via measures of glomerular layer immunofluorescence and whole-bulb protein expression (Parrish-Aungst *et al*., 2011). Decreased expression of *Gad1* mRNA and GAD67 protein have also been described following chemical ablation of olfactory sensory neurons (Lau & Murthy, 2012) or unilateral naris occlusion (Banerjee *et al*., 2013; Wang *et al*., 2017). However, these studies used longer-duration manipulations and did not distinguish between different bulbar neuronal populations. Therefore, either 1-3 days of unilateral naris occlusion is insufficient to induce activity dependent changes in GAD67, or measures based on immunofluorescence intensity may not be sensitive enough to always detect changes in GAD67, or these changes are driven in neurons not labelled in the DAT-tdTomato mouse line.

### Experience-dependent plasticity of dopamine-synthesising enzymes

We report here the first evidence for an activity-dependent change in olfactory bulb DDC levels. This effect was very small and only transient, with relative DDC immunofluorescence in DAT-tdTomato neurons decreasing slightly after 1 day of naris occlusion and returning back to baseline after just 3 days of sensory deprivation. It was also in the opposite direction to the trend we observed in whole-bulb qPCR data, which suggested a transient *increase* in *Ddc* mRNA after 1 day occlusion. Perhaps this mRNA increase reflects a transcriptional rebound already in operation after just 24 h deprivation, driving DDC protein rapidly back towards control levels. The small and temporary experience-dependent DDC immunofluorescence alteration we observed might explain why previous studies of potential bulbar DDC plasticity – which used much longer periods of sensory manipulation - did not find similar effects (Stone *et al*., 1990; Cave *et al*., 2010). However, expression of DDC in the midbrain has been reported to be affected within a few hours of activation or blockade of dopamine receptors (Hadjiconstantinou *et al*., 1993; Zhu *et al*., 1993). Further work is needed to confirm whether downregulation of bulbar DDC after 1 day of occlusion is a consistent activity-dependent modification. If so, it might reflect an initial strategy whereby dopaminergic neurons downregulate multiple nodes in the dopamine synthesis pathway, before sufficiently decreased TH activity can alone – as the rate-limiting step – control levels of transmitter production.

Our data showing decreased TH expression after olfactory sensory deprivation are entirely consistent with a large body of prior work describing this effect (Nadi *et al*., 1981; Baker *et al*., 1983, 1984, 1993, 1999; Kosaka *et al*., 1987; Baker, 1990; Stone *et al*., 1990; Cho *et al*., 1996; Philpot *et al*., 1998; Saino-Saito *et al*., 2004; Bastien-Dionne *et al*., 2010; Cave *et al*., 2010; Weiss *et al*., 2011; Grier *et al*., 2016), including some indications that it can occur after relatively brief manipulations (Cho *et al*., 1996; Akiba *et al*., 2007; Chand *et al*., 2015; Galliano et al., 2021). It is novel, however, that our methodological approach demonstrated activity-dependent TH alterations in cells identified in an entirely TH-independent manner. As discussed above, this approach was not completely selective, labelling a minority of calretinin-positive neurons in addition to dopaminergic cells. However, since we never observed a DAT-tdTomato neuron that was both calretinin-positive and dopaminergic, and since the proportions of calretinin-positive cells did not change significantly with occlusion (Fig. 3), this off-target labelled sub-population cannot account for the relative changes in TH and DDC immunofluorescence we observed. Indeed, if anything, the presence of a significant proportion of essentially TH- or DDC-negative cells amongst the DAT-tdTomato samples we analysed may have made our ability to detect relative differences in enzyme levels less sensitive.

We also note the caveat that, of course, changes in immunofluorescence intensity cannot be used to infer activity-dependent plasticity in *absolute* levels of protein expression. With this approach, results can be easily confounded by variability in the duration of immunohistochemistry steps, the composition of reagents, and/or individual image acquisition settings. We did take every possible measure to mitigate against such factors, comparing tissue from mice that were fixed on the same day with the same fixative solution, comparing sections that were physically co-embedded prior to all stages of histological processing, and comparing images that were taken on the same day with the same acquisition settings. We also ensured that we only ever made *relative* comparisons of immunofluorescence. So, while we are confident that, for example, TH levels decreased in DAT-tdTomato cells after 1 day of sensory deprivation, we do not claim any insight into the magnitude of this decrease. We do, however, have corroborative, fully quantitative qPCR data to show that – at least at the whole-bulb level – brief sensory deprivation is associated with an approximate halving of *Th* mRNA levels.

### Mechanisms

It remains unclear precisely how activity-dependent changes in TH expression occur, though the immediate depolarisation-activated pathways appear to require signalling through L-type CaV1 Channels and cAMP (Cigola *et al*., 1998), and *Th* gene expression is at least partly controlled by changes in secondary DNA structure influencing the binding of heterogeneous nuclear ribonucleoproteins (hnRNPs) to its proximal promoter (Banerjee *et al*., 2014; Wang *et al*., 2017). Our findings suggest that these processes can work over reasonably quick timescales. However, given the similarity in promoter-based changes in secondary structure and hnRNP binding between the *Th* and *Gad1* genes (Wang et al., 2017), it will be interesting to consider how these processes can work more rapidly, more specifically, and/or be coupled with targeted changes in mRNA and protein turnover, to produce selective downregulation of TH without any changes in GAD67 after a few days of naris occlusion.

### Implications for sensory processing

Plasticity of TH expression is believed to be a way in which the olfactory bulb can tune its levels of dopaminergic signalling according to changes in incoming sensory-evoked activity. Given some of the known effects of dopamine in glomerular networks, this could potentially have the effect of counteracting input activity perturbation. For example, the best-described circuit-level effect of dopamine in the bulb is its inhibitory action on the release of glutamate from olfactory sensory neuron axon terminals, which it achieves via D2 receptor activation (Hsia *et al*., 1999; Berkowicz & Trombley, 2000; Ennis *et al*., 2001; Maher & Westbrook, 2008; McGann, 2013; Vaaga *et al*., 2017). If sensory input levels are chronically decreased, it might be adaptive to reduce the strength of this inhibition, perhaps by decreasing expression of the rate-limiting enzyme for neurotransmitter synthesis. This might put the system into a state of higher-gain readiness for the low levels of input that will initially arrive if and when sensory stimulation recommences. Indeed, the volume-transmission nature of sensory neuron presynaptic inhibition (Pinching & Powell, 1971; McGann, 2013), coupled with the high threshold and slow dynamics of bulbar dopamine release (Borisovska *et al*., 2013), might make for a slow modulatory process that is well suited to days-scale modification via plasticity of enzyme expression. However, given the strong presynaptic inhibition also mediated by GABA_B_ receptor activation on OSN terminals (McGann, 2013), it is unclear why dopamine-but not GABA-synthesising enzymes are targets for rapid activity-dependent regulation in this circuit. It may be that GABAergic inhibition is modulated by distinct plastic mechanisms that do not target transmitter synthesis. Alternatively, distinct mechanisms of GABAergic and dopaminergic inhibition that can occur within the same presynaptic terminals (Burke *et al*., 2018) might be individually tuned according to sensory experience.

In contrast, the potential functional implications of TH plasticity on the less-understood postsynaptic effects of bulbar dopamine transmission are harder to predict. Dopamine can act via D1-type receptors to promote rebound firing after interglomerular GABAergic inhibition of external tufted cells (Liu *et al*., 2013), so perhaps decreased TH levels might lead to more prolonged lateral inhibition within the glomerular layer. The implications of such an effect for information processing in sensory-deprived bulbar circuits might be best addressed via computational network modelling. Most important, though, is the fact that, although experience-dependent plasticity in olfactory bulb TH expression has been described for decades, it is still not known how (or whether) changes in TH mRNA and protein levels give rise to alterations in the amount of dopamine actually released. Future experiments should be able to control the activity of dopaminergic neurons whilst monitoring the resulting effects of dopamine release (e.g. Vaaga *et al*., 2017), in order to quantitatively compare the outcome in control versus sensory-deprived tissue.

## Author Contributions

DJB, ML & MG designed experiments; DJB, AC & ML performed experiments; DJB, ML & MG analysed data; DJB, ML & MG wrote the paper.

## Acknowledgements

This work was supported by a Medical Research Council 4-year PhD studentship to DJB, and a Leverhulme Trust Research Grant (RPG-2016-095) and ERC Consolidator Grant (725729; FUNCOPLAN) to MSG. We wish to thank Beatriz Rico for access to equipment, Laura Andreae and Elisa Galliano for comments on the manuscript, and Elisa Galliano, Christiane Hahn and other past and present members of the Grubb laboratory for technical assistance and stimulating discussion.

## Data accessibility

The data that support the findings of this study will be fully and openly available upon publication in a public repository that issues datasets with DOIs.

